# Genetic variability and vector competence of *Aedes aegypti* populations from Kisumu and Busia Counties, Western Kenya, for Chikungunya and Zika viruses

**DOI:** 10.1101/2023.07.14.549035

**Authors:** Victor O. Anyango, Solomon Langat, Francis Mulwa, James Mutisya, Hellen Koka, Collins Okoyo, Edith Chepkorir, Samson Konongoi, Anncarol Karanja, Glennah Kerubo, Rosemary Sang, Joel Lutomiah

## Abstract

*Aedes aegypti* is the primary vector of several arboviruses, including dengue virus (DENV), chikungunya virus (CHIKV), yellow fever virus (YFV), and Zika virus (ZIKV). This vector is widespread globally in tropical and subtropical areas, but also found in temperate areas. Kenya experienced its first chikungunya outbreaks in Lamu County in 2004 and later in Mandera: 2016, and Mombasa: 2017. While there is yet to be a report of Zika outbreaks in Kenya, sero-surveillance studies indicate low-level transmission of this virus in coastal and northern parts of the country. Despite the presence of *Ae. aegypti* in Kisumu and Busia counties in sufficient densities, and free movement of people between the coast and the two western Kenya counties, no outbreaks of either disease have been reported in these regions. To investigate this phenomenon, we collected *Ae. aegypti* mosquitoes from county headquarter towns near railway stations connecting the coast and western Kenya and reared them under controlled laboratory conditions. The mosquitoes were then assessed for genetic variability using CO1 genes as well as their efficiency to transmit viruses using Laboratory colonies (F_1_) of the field mosquitoes challenged with an infectious blood meal containing CHIKV and ZIKV.

Genetic analysis revealed the presence of both *Ae. aegypti* subspecies, (*Ae. aegypti aegypti* [*Aaa*] and *Ae. aegypti formosus* [*Aaf*]) in the two western Kenya counties, with *Aaf* being dominant (19:8 for Kisumu samples and 25:6 for Busia samples). Additionally, pairwise comparison revealed minimal genetic differentiation (0.62%) between the study populations, with a high genetic variation (99.38%) observed within each population, indicating significant diversity within individual populations. *Ae. aegypti* populations from Kisumu and Busia counties exhibited competence for CHIKV, with infection, dissemination, and transmission rates of 55.2%, 85.5%, and 27.1% for Kisumu; and 57.8%, 71.8%, and 25% for Busia populations, respectively. There was no significant difference in vector competence between these two populations. Interestingly, neither population was competent for ZIKV. In conclusion, the data shows that the *Ae. aegypti* populations in the two cities were homogeneous. This could explain the observed similarity in vector competence for CHIKV and ZIKV.

**Author Summary:** Our study investigated the genetic variability and vector competence of *Ae. aegypti* mosquito populations in Kisumu and Busia Counties to CHIKV and ZIKV; revealing the presence and even distribution of both *Aaa* and *Aaf* subspecies. We also found that the *Ae. aegypti* populations from the two counties were not genetically differentiated. Furthermore, our study revealed that the *Ae. aegypti* mosquitoes from Kisumu and Busia counties were competent for CHIKV but may be refractory to ZIKV infection. These findings highlight the importance of continued monitoring of *Ae. aegypti* populations and their potential for arboviral disease transmission in the region.

## Introduction

*Ae. aegypti* is the primary vector for Zika and Chikungunya viruses and is believed to have originated from Africa (1). Originally, *Ae. aegypti* was a single species inhabiting the African forests, breeding in tree holes, rock pools, and still water, and feeding on wild animals (2). Later genetic analyses identified two subspecies of *Ae. aegypti*: *Ae. aegypti aegypti* (*Aaa)* and *Ae. aegypti formosus* (*Aaf*), which are morphologically, behaviorally, and genetically differentiated (2,3). *Aaf* is darker in color, zoophilic and confined to forests, while *Aaa* is lighter, anthropophilic/ prefers human blood and breed in artificial habitats, thus is also referred to as the domestic sub species (2,4). Research suggests that *Aaf* may have been brought to the New World aboard ships during the slave trade, where it evolved into *Aaa*, a more proficient carrier of arboviruses with a specialization for biting humans (3,5). Consequently, *Aaa* subspecies, is thought to have been introduced to the African continent via the continent’s seaports (2,6) and there may be a possibility that they migrated to other parts via the available modes of transport which was mainly the railway line. The frequent outbreaks of arboviruses, such as chikungunya and dengue fever, along Africa’s coastal areas lends credence of the hypothesis that *Aaa* may have been reintroduced to Africa’s coast (7). Moreover, genetic analysis of *Ae. aegypti* mosquitoes has revealed that the sylvan form of *Ae. aegypti* is more common in Africa, while the domestic form is more prevalent in Europe, South and North America and Asia (2,4).

*Ae. aegypti aegypti* (*Aaa*) has been shown to be more efficient in transmitting these viruses than *Aaf,* due to its preference for human blood meal, and ability to feed multiple times during a single gonotrophic cycle (8–10). Furthermore, Aubry *et al.* (2020) demonstrated that the *Aaf* (4) are less susceptible to ZIKV infection compared to the domestic forms, *Aaa*. Additionally, Grard *et al.* (2014) demonstrated that the 2007 Zika virus disease outbreak in Gabon was driven by *Ae. albopictus* despite the presence of *Ae. aegypti* mosquitoes in the country (11,12), probably because this species is predominantly *Aaf* (12). Interestingly, the 2015 – 2016 Zika virus disease outbreaks in Brazil and the Caribbean were linked to the domestic form, *Aaa* (13). These findings highlight the significant variability in susceptibility and vector competence between the two subspecies of *Ae. aegypti*, underscoring the complex dynamics of arbovirus transmission.

First isolated from the febrile rhesus macaque monkey in Uganda’s Zika Forest in 1947 (14), ZIKV is primarily transmitted through bites of an infected *Ae. aegypti* mosquito among other modes (13–15). It causes Zika virus disease which is often associated with mild and self-limiting febrile illness with symptoms lasting 2 to 7 days. However, in extreme cases, it may result in neurological disorders such as microcephaly in newborn babies as well as Guillain-Barre Syndrome, myelitis, and neuropathy among infected adults and teenagers, making it a significant public health concern (15–18). Zika virus disease outbreaks have been reported in the Americas, Europe, Asia and the pacific, all of which are considered endemic zones for the virus (19–21). Although Kenya harbors the primary vector of ZIKV, no cases of the zika virus disease have ever been reported in the country. However, the detection of ZIKV antibodies in northern, western and coastal Kenya (22,23) indicate the possible widespread but low-level circulation of the virus among populations in the country.

Similar to ZIKV, CHIKV was first identified in Africa, specifically in Tanzania in 1952(24). Since then, it has spread to more than 110 countries across different continents, including Africa, Asia, the Indian subcontinent, Europe, and the Americas (25–30). CHIKV is primarily transmitted through bites of infected mosquitoes particularly of the *Ae.* species and is often characterized by self-limiting symptoms including febrile illnesses and severe arthritis. Outbreaks of CHIKV have been reported in several regions of Kenya, including Lamu in 2004 and 2021-2022 (26,29), Mandera in 2016 (28) as well as in Mombasa in 2017 and 2018 (31). Moreover, It is estimated that approximately150 people in Kenya have died due to Chikungunya fever, (26,32) making the virus a significant public health concern.

Although studies have shown the presence of *Ae. aegypti* mosquitoes across Kenya, no outbreaks have been reported in western Kenya (7). This suggests that the transmission and impact of both viruses in this region have been inconsequential. However, the widespread distribution of the vector and detection of virus antibodies in this region indicate a potential risk transmission (7,23). Therefore, the aim of this study was to examine the entomological factors that could potentially explain the absence of reported outbreaks in western Kenya. Specifically, the genetic variability and vector competence of *Ae. aegypti* populations collected from two western Kenya counties: Kisumu and Busia was analyzed to determine the genotypes present in the region as well as to understand the mosquitoes’ ability to acquire and transmit the viruses.

## Results

### Genetic variability of *Ae. Aegypti* populations from Kisumu and Busia Counties

DNA sequences were analyzed, and a diversity table generated. 27 DNA sequences were obtained from the mosquitoes collected from Kisumu, and 31 from mosquitoes collected from Busia. The DNA analysis generated 18 haplotypes for the Kisumu population and 23 for the Busia population. The haplotype diversity of mosquitoes from Kisumu and Busia was 0.96011 and 0.93763 respectively, while the nucleotide diversity was 0.00915 for both Kisumu and Busia populations, and AMOVA, 0.00575 (**Table 1**).

**Table 1:** Diversity table. Genetic variability of *Ae aegypti* mosquitoes from Kisumu and Busia Counties.

| Sites | n | Hn | Hd | $\pi$ |
| --- | --- | --- | --- | --- |
| Kisumu | 27 | 18 | 0.96011 | 0.00913 |
| Malaba | 31 | 23 | 0.93763 | 0.00757 |
| Total | 58 | 37 | 0.96128 | 0.00832 |

### Pairwise comparison of *Ae. Aegypti* sequences from Kisumu and Busia counties

The pairwise comparisons conducted to determine the study populations’ variation indexes showed that, percentage variation among the two study populations was 0.62%, with 1 df, while variation within individual populations was 99.38%, with 56 degrees of freedom (**Table 2**).

**Table 2:** Pairwise comparison table of *Ae. Aegypti* sequences from Kisumu and Busia counties.

| Source of variation | df | Sum of squares | Variance components | Percentage of variation |
| --- | --- | --- | --- | --- |
| Among populations | 1 | 3.73 | 0.01983 Va | 0.62 |
| Within populations | 56 | 176.822 | 3.15754 Vb | 99.38 |
| **Total** | **57** | **180.552** | **3.17736** |  |

df degrees of freedom

### Phylogenetic analysis

Phylogenetic analysis of the 18 and 23 haplotypes obtained from the Kisumu and Busia mosquito sequences showed the existence of the two *Ae. Aegypti* sub species: *Aaa* and *Aaf* in both sites (**Fig 1**). *Aaf* were found to be more abundant in both sites compared to *Aaa*. Based on the phylogenetic analysis, there were no geographical clustering patterns of the sequences from either site, clustering was based on the sub species (*Aaa* and *Aaf*) as opposed to the study sites. Overall, there was high bootstrap support for the haplotype clustering which was above 50%.

**Fig 1:**
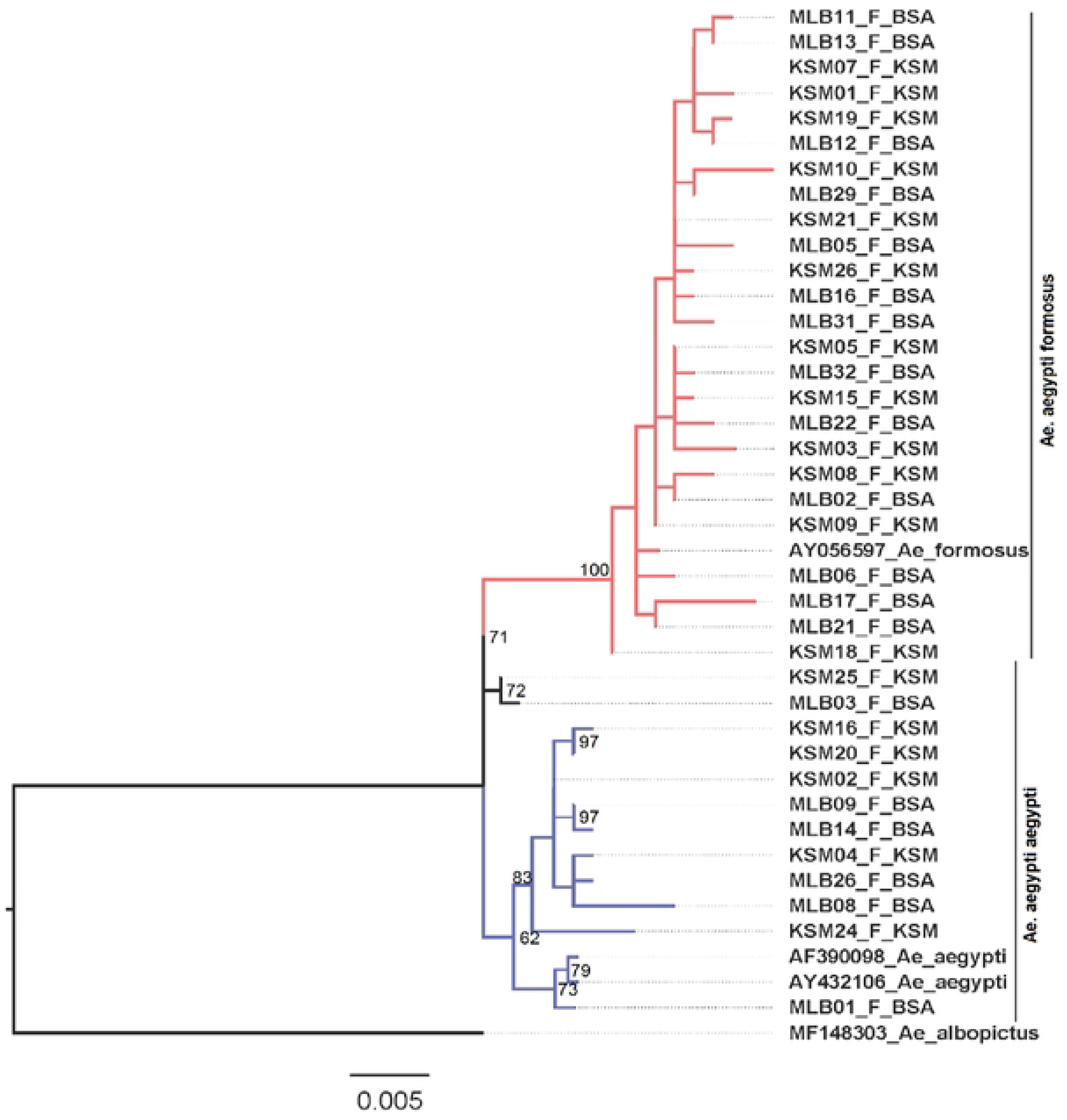
Maximum likelihood tree showing the genetic variability of *Ae. Aegypti* species collected from Kisumu and Busia Counties. The red and black colored branches represent haplotype sequences from the two sites clustered towards *the Aaf* genotype, while the blue colored branches represent those that clustered towards *the Aaa* genotype (the distinction has also been made by the two vertical lines on the far right of the diagram). The bootstrap supporting haplotypes that clustered towards the Aaf was 72% while that which supported haplotypes that clustered towards *Aaa was* 62%.

### Vector competence of *Ae. aegypti* mosquitoes from Kisumu and Busia for CHIKV and ZIKV

A total of 260 and 192 mosquitoes from the two study sites were challenged with Log 10^6.68^ PFU/ml of CHIKV and Log 10^4.52^ PFU/ml ZIKV infectious blood meals respectively. Overall, approximately 56.5% of all mosquitoes challenged with CHIKV were susceptible, 78.2% disseminated the virus and 26.1% of those that disseminated the virus showed transmission capabilities as was seen by the presence of CHIKV virus in their saliva.

For mosquito populations from Kisumu, a total of 125 mosquitoes were challenged with CHIKV infectious blood meal. Approximately 55.2% of the mosquitoes challenged were infected, 85.5% of the infected mosquitoes disseminated the virus and 27.1% exhibited transmission capabilities. On the other hand, for mosquitoes from Busia, a total of 135 mosquitoes were challenged with CHIKV infectious blood meal. Approximately 57.8% of them were infected, 71.8% of the infected mosquitoes disseminated the virus and 25.0% exhibited transmission capabilities. The proportional test of differences showed no significant difference between infection, dissemination, and transmission rates of mosquito populations from Kisumu and Busia challenged with CHIKV. Interestingly, mosquito populations from Kisumu and Busia counties that were challenged with ZIKV infectious blood meals were not susceptible to either virus. A total of 192 mosquitoes from Kisumu and Busia were challenged with ZIKV infectious blood meal and their body homogenates assesses for virus particles by inoculation in Vero cells. None of the inoculum showed CPE therefore, no further assays were conducted. (**Table 3**).

**Table 3:** Vector competence of *Ae. aegypti* mosquitoes from Kisumu and Busia counties for CHIKV and ZIKV.

| Site | The proportion of mosquitoes infected with CHIKV.<br>% (95%CI); n=260 |  |  | The proportion of mosquitoes infected with ZIKV.<br>% (95%CI); n=192 |  |  |
| --- | --- | --- | --- | --- | --- | --- |
|  | Body<br>(Infection) | Leg<br>(Dissemination) | Saliva<br>(Transmission) | Body | Leg | Saliva |
| Overall | 56.5 (50.8-62.9); n=147 | 78.2 (71.8-85.2); n=115 | 26.1 (19.2-35.5); n=30 | 0 | 0 | 0 |
| Site |  |  |  |  |  |  |
| Busia<br>(n=135) | 57.8 (50.0-66.7); n=78 | 71.8 (62.5- 82.5); n=56 | 25.0 (15.9-39.4); n=14 | 0 | 0 | 0 |
| Kisumu<br>(n=125) | 55.2 (47.1-64.6); n=69 | 85.5 (77.5-94.2); n=59 | 27.1 (17.8-41.2); n=16 | 0 | 0 | 0 |
| Proportion test of the difference (diff, *p value*) | Diff=0.026, p=0.751 | Diff=-0.137, p=0.072 | Diff=-0.021, p=0.896 | - | - | - |

**n: No. of mosquitoes, CI: confidence interval, p. Probability value**

The study further compared rates of infection, dissemination, and transmission based on the virus incubation period in the sample mosquitoes. For Kisumu, approximately 44.2% of the mosquitoes processed after 5 days of incubation were infected with CHIKV. Of those infected, 78.9% disseminated the virus, and 33.3% of those that disseminated the virus showed transmission capabilities. At 10 dpi, approximately 60.9% of the processed mosquitoes were infected with CHIKV. Of those infected, 89.3% disseminated the virus, and 40% of those with disseminated infection showed transmission capabilities. Similarly, at 14 dpi, approximately 61.1% of the processed mosquitoes were infected with CHIKV, and 86.4% of those infected disseminated the virus. However, only 5.3% of those that disseminated the virus showed transmission capabilities by the presence of CHIKV in their saliva. (**Table 4**).

**Table 4:** Vector competence *Ae aegypti* mosquitoes from Kisumu and Busia County to CHIKV based on incubation periods.

|  | Kisumu |  |  |  |  | Busia |  |  |  |  |
| --- | --- | --- | --- | --- | --- | --- | --- | --- | --- | --- |
| Day | Day 5 | Day 10 | Day 14 | Proportion test of difference at day 10 | Proportion test of difference at day 14 | Day 5 | Day 10 | Day 14 | Proportion test of difference at day 10 | Proportion test of difference at day 14 |
| Chikungunya |  |  |  |  |  |  |  |  |  |  |
| Body | 44.2<br>(31.6-61.8);<br>n=19/43 | 60.9<br>(48.3-76.7);<br>n=28/46 | 61.1<br>(47.1-79.3);<br>n=22/36 | Diff=0.167,<br>p=0.259 | Diff=0.169,<br>p=0.280 | 59.6<br>(47.1-75.4);<br>n=28/47 | 48.9<br>(36.3-65.9);<br>n=22/45 | 65.1<br>(52.3-81.0);<br>n=28/43 | Diff=0.107,<br>p=0.450 | Diff=-0.055,<br>p=0.671 |
| Leg | 78.9<br>(62.6-99.6);<br>n=15/19 | 89.3<br>(78.5-101.5);<br>n=25/28 | 86.4<br>(73.2-102);<br>n=19/22 | Diff=0.104,<br>p=0.367 | Diff=0.075,<br>p=0.562 | 62.1<br>(46.7-82.5);<br>n=18/27 | 81.8<br>(67.2-99.6);<br>n=18/22 | 74.1<br>(59.3-92.6);<br>n=20/27 | Diff=0.197,<br>p=0.199 | Diff=-0.12,<br>p=0.427 |
| Saliva | 33.3<br>(16.3-68.2);<br>n=5/15 | 40.0<br>(24.7-64.6);<br>n=10/25 | 5.3 (7.8-35.5);<br>n=1/19 | Diff=0.067<br>p=0.801 | Diff=0.279,<br>p=0.572 | 44.4<br>(26.5-74.5);<br>n=8/18 | 16.7<br>(59.3-46.8);<br>n=3/18 | 15.0<br>(52.8-42.6);<br>n=3/20 | Diff=0.277,<br>p=0.396 | Diff=0.294,<br>p=0.367 |

For mosquito populations from Busia County,59.6% of those challenged with CHIKV were infected at 5dpi. Of the infected mosquitoes, 62.1% disseminated the virus, and 44.4% of those that disseminated the virus showed transmission capabilities. The infection rate decreased from 59.6% at 5 dpi to 48.9% at 10 dpi, while the dissemination rate increased from 62.1% to 81.8%. However, the proportion of mosquitoes that showed transmission capabilities decreased significantly to 16.7% given the high number of mosquitoes that disseminated the virus. At 14 dpi, the infection dissemination and transmission rates were 65.1%, 74.1% and 15% respectively (Table 4).

The results of the plaque assay performed on positive mosquito bodies, legs and saliva from both the Kisumu and Busia samples confirmed the presence of live viral particles, with mean titers of 10^5.916^ and 10^4.165^ for Kisumu and 10^5.8325^ and 10^3.99^ for Busia, respectively (Table 5). These findings indicate that the cell culture results above were true. However, no plaques were observed in any of the saliva samples.

**Table 5:** Mean titers of mosquito bodies and legs titers.

|  | Mean body titers (PFU) | Mean Leg titers (PFU) |
| --- | --- | --- |
| Kisumu | $10^{5.916}$ | $10^{4.165}$ |
| Busia | $10^{5.8325}$ | $10^{3.99}$ |

## Discussion

In this study, we determined the genetic variability and vector competence of *Ae. aegypti* populations from Kisumu and Busia counties, Kenya for CHIKV and ZIKV. The findings presented herein provide critical insights into the transmission dynamics of these viruses.

Our study observed a high haplotype diversity (0.96011 and 0.93763) and low nucleotide diversity (0.00913 and 0.00757) in mosquito populations from Kisumu and Busia counties respectively. One possible explanation for this is the introduction of similar populations into the region. Additionally, the low nucleotide diversity observed suggests limited-to-no-change in the genetic materials of *Ae. aegypti* populations. Pairwise comparison of *Ae. aegypti* populations from Kisumu and Busia counties showed a higher variation within the populations in each site and low variation between the two sites. Based on these findings, we conclude that there is no significant genetic variability between *Ae. aegypti* populations from the Kisumu and Busia Counties. However, the variation within each population is significant, something that is worth investigating.

The phylogenetic analysis data revealed the existence of both *Aaa* and *Aaf* sub-species in the cities, with *Aaf* being dominant (19: 8 for *Aaf : Aaa* in Kisumu and 25:6 for *Aaf : Aaa* in Busia). These findings are in tandem with a previous study by Futami *et al*. (2020) which assessed the geographical distribution of *Aaa* and *Aaf* in Kenya (7). The study found that *Aaa* was more prevalent in the coastal region, with decreasing abundance as distance from the sea increased towards western Kenya where higher *Aaf* abundance was recorded. Additionally, a study by Agha *et al*. (2022) that included phylogenetic analysis of *Ae. aegypti* mosquitoes from Kisumu showed the existence of both *Aaa* and *Aaf* in the region, providing further support for these findings (33). Collectively, these results suggest that the distribution and abundance of *Aaa* and *Aaf* sub-species in western Kenya may have implications for vector competence and disease transmission in the region, since *Aaa* has been documented as an efficient transmitter of arboviruses due its anthropophilic behavior (2) while *Aaf* is zoophilic. Therefore, our findings highlight the importance of continued surveillance and monitoring of mosquito population dynamics and virus circulation in the region for early detection, outbreak prevention and mosquito control.

Our study also examined the vector competence index of *Ae. aegypti* mosquitoes from the two study sites. The results of our study revealed that, *Ae. aegypti* populations from Kisumu and Busia counties were competent for transmitting the CHIKV. Despite the average susceptibility to CHIKV recorded in the study, a huge proportion of mosquitoes that were infected were able to disseminate the virus and this cut across the two study sites. Interestingly, the number of mosquitoes with successful transmission reduced, as only a small proportion of mosquitoes with disseminated infection transmitted the virus. The role of midgut infection (MIB) and escapes (MEB) barriers as well as salivary gland infection (SGI) and escape (SGEB) barriers in explaining this phenomenon cannot be overlooked. The average infection rate observed in this study is an indication of a well-established and strong MIB in mosquitoes that failed to establish CHIKV infection. These barriers, created by the midgut epithelial cells, peritrophic matrix and midgut microbiota have been associated with reduction in vector viremia as they limit/prevent the establishment of pathogens within the mosquito’s body (34,35). Moreover, of all the susceptible mosquitoes, 78.2% disseminated the virus, an indication of a weakened MIB and MEB among this population. Conversely, only 26.1% of the mosquitoes that disseminated the virus demonstrated transmission capabilities, suggesting an overall strong SGIB and SGEB among the populations. Salivary gland infection and escape barriers are responsible for preventing the establishment of pathogen in mosquitos’ salivary glands as well as the shedding of the pathogen into the mosquito saliva for transmission (35). Therefore, the study concludes that MIB and MEB as well as the salivary gland barriers are key factors that may affect vector competence among mosquitoes from Kisumu and Busia counties and needs further investigation.

However, it is important to note that the forced salivation procedure used in this study has been documented as a less efficient method for determining mosquito transmission abilities, given its propensity to underestimate virus transmission (36). Therefore, the presence of virus particles in mosquito legs is a more reliable predictor of its transmission capabilities than forced salivation methods (36). The findings of this study may underestimate the true extent of virus transmission, as transmission levels in natural settings with vertebrate hosts present could be higher compared to laboratory settings that utilize artificial capillaries.

A study by Agha *et al.,* (2017) demonstrated that, *Ae. aegypti* populations from Kisumu were competent to CHIKV further supporting the findings of this study (33). Our study also demonstrated that CHIKV-infected mosquitoes could transmit the virus as early as 5 dpi, an indication of the study mosquito’s ability to efficiently transmit the virus within a short period of time, posing a substantial risk for the rapid spread of the virus within a population. These findings align with previous studies that have also observed a shorter incubation period of CHIKV in mosquito vectors (33,37).

On the contrary, mosquito species challenged with ZIKV were not susceptible, indicating that populations of *Ae. aegypti* from western Kenya may exhibit refractoriness to ZIKV. The results further emphasize the importance of MIB and mosquito microbiota in protecting *Ae .aegypti* from Western Kenya against ZIKV infection. An increasing body of evidence suggests that mosquito microbiota may have a protective effect against viral infections (38). Therefore, further investigation is needed to explore the potential role of mosquito microbiota in elucidating the aforementioned phenomena.

These results also confirm the role of geographic isolation to variation in vector competence of mosquito populations to arboviruses(39) given the closeness of western Kenya, especially Busia, to the zika Forest in Uganda where ZIKV was first discovered,

In conclusion, our findings demonstrate the presence of *Ae. aegypti* subspecies (*Aaa* & *Aaf*) in Western Kenya. We further demonstrate that, *Ae. aegypti* populations from the two study sites were not genetically different, although variation within each population were found to be high indicating significant diversity within individual populations. Our study also showed that *Ae. aegypti* populations from Kisumu and Busia counties were competent for CHIKV but refractory to ZIKV. Based on these findings, we recommend that future studies consider using more robust SNPs analysis to determine the genetic heterogeneity of *Ae. aegypti* in these regions. We also recommend that further studies be conducted using higher ZIKV titers and different ZIKV strains to corroborate the findings of this study. In addition, we recommend that studies be conducted to explore the role of *Ae. aegypti* microbiota in protecting mosquitoes against ZIKV and the mechanisms involved.

## Materials and Methods

### Study sites

The study was conducted in two western Kenya counties: Kisumu and Busia (**Fig 2**). *Ae. Aegypti* mosquitoes were collected in county headquarter towns within the vicinity of the old railway stations old railway stations and transferred to the Kenya Medical Research Institute (KEMRI) Biosafety Level II insectary. Kisumu and Busia have a population of approximately 1,155,574 and 893,681 respectively according to the Kenyan population census of 2019 (40). The climate comprises a warm and humid tropical climate with temperatures ranging from 22°C to 30°C for Kisumu County and 18°C to 34°C for Busia County throughout the year making the two counties conducive habitats for mosquitoes.

**Fig 2:**
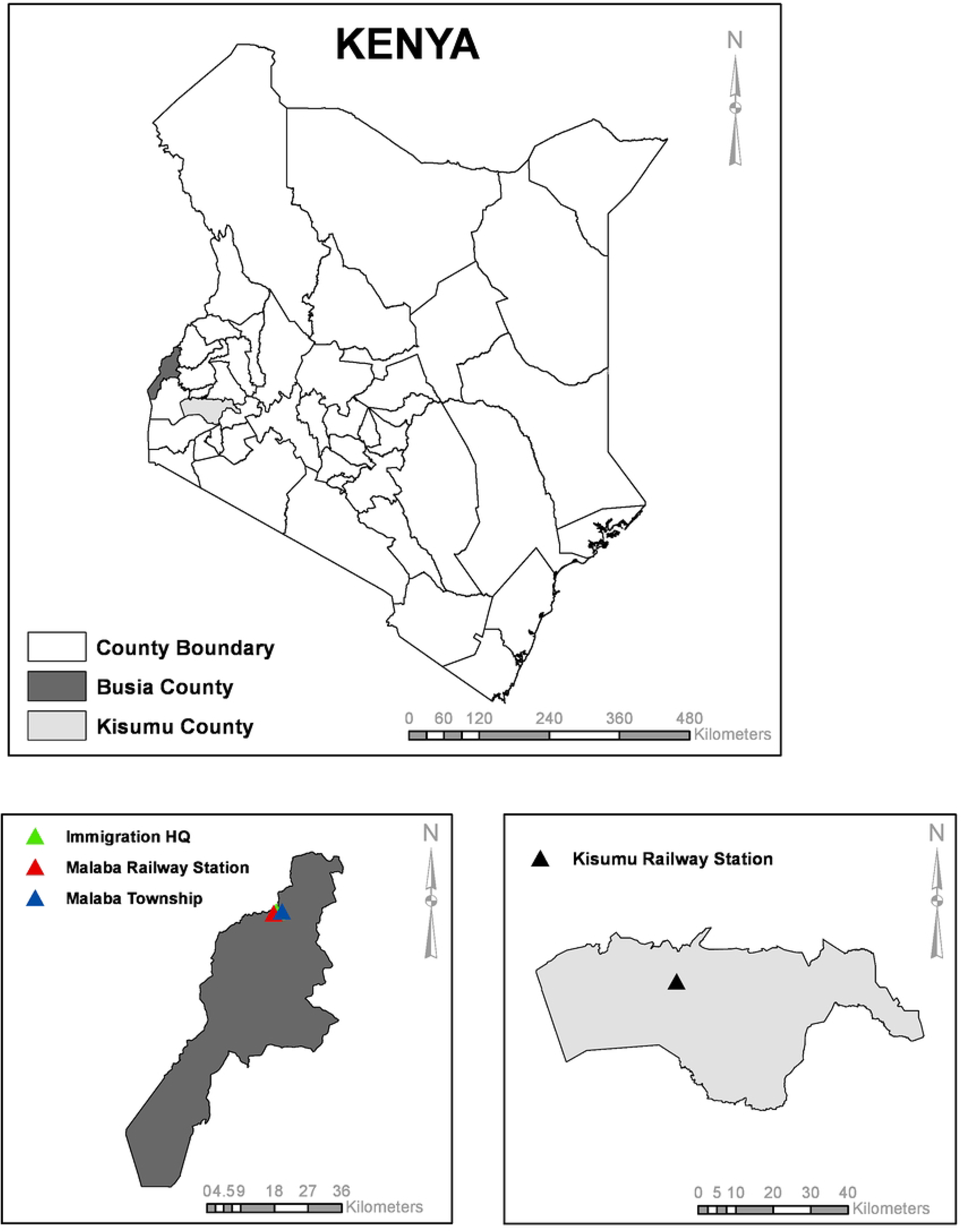
Map of Kenya showing the two study sites (Kisumu and Busia counties)

### Mosquito collection and rearing

From June to September 2021, *Ae. aegypti* mosquitoes were collected in the form of eggs, larvae, pupae, and adults. Adult mosquitoes were collected using carbon baited BG sentinel traps during the day as well as at night. For the eggs collection oviposition cups (ovitraps) lined with oviposition papers and half-filled with water were used (41). The ovitraps were placed under trees, bushes, residential and commercial areas, and old vacated railway buildings, for three days to allow mosquitoes to lay eggs. In addition, the immature stages of the mosquito: larvae and pupae were collected from various natural and artificial habitats such as tree holes, old tires, concrete holes, stagnant water, abandoned cans, and abandoned water tanks using scoopers and teat pipettes, and transferred into Whirl packs. Both the adult and immature mosquitoes were transported to the biosafety level II insectary lab at KEMRI in Nairobi, Kenya, for further analysis.

The Identified F_0_ *Ae. aegypti* were reared under controlled insectary conditions (28 ± 1°C, 12:12 hour light-dark cycle, and 75 ± 5% relative humidity) with access to 10% glucose (42). The field-collected eggs were hatched in a 24 × 34 × 9 cm tray half filled with dechlorinated water, larvae were maintained using standard Tetramin (Tetra) fish food and monitored daily for the presence of pupa. The pupa stages were picked from the tray and transferred to a 200ml beaker containing clean, dechlorinated water, which was then placed in an insect cage (BugDorm) measuring 30 x 30 x 30 cm and allowed to emerge into adults (42). Approximately 600 pupae were placed in single cages (42). Adult mosquitoes that emerged from the developmental stages collected from the field were identified under a dissecting microscope and *Ae. aegypti* species isolated. The F_0_ adult mosquitos were blood-fed, and F_1_ eggs collected for subsequent vector competence assays.

### Assessment of genetic variability Ae. Aegypti DNA extraction

Approximately 32 field collected female mosquitoes (F_0_) were selected from each county (Kisumu and Busia) by sampling from various collection sites and collection methods. Individual mosquito legs were then isolated using sterile forceps and placed in a sterile Rnase/Dnase-free Eppendorf tube containing 3 pieces of 2.0mm Zicorna beads. 200ml of Phosphate buffered saline was then added into each Eppendorf tube and the sample homogenized in an Omni Bead Rapture 24 (OMNI International) set to run at the speed of 2.10mm for 20 seconds. *Quick-*DNA*™* Miniprep Kit (ZYMO RSEARCH) (43) was then used to extract DNA from the homogenates following the manufacturers directives

### Polymerase chain reaction

Polymerase Chain Reaction (PCR) was used to amplify the target mtDNA (COI) in mosquito samples using the universal reverse primer HCO2198 and forward primer LCO1490 (44). The PCR master mix was prepared in a Biosafety level II molecular laboratory and contained 12.5µl Amplitaq solution, 1µl forward primer, 1µl reverse primer, and 8.5µl sterile nuclease free water. Next, 23µl of the master mix was transferred into well-labelled RNase-free, 0.2 ml (8-strip format, Invitrogen) PCR tubes and 2µl of sample DNA added into each tube. The tubes were tightly sealed, vortexed, and placed in a Thermocycler set to run 40 reaction cycles of 92°C for the 30s, 43–52°C for 30s, and 72°C for 60s (29). The amplicons were visualized in a 2% agarose gel with an expected band size of approxiately 650bp, stained with ethidium bromide and sent to macrogen company for sequencing (45).

### Determination of vector competence of *Ae. aegypti* to ZIKV and CHIKV

#### Virus amplification

Chikungunya virus strain isolated during the 2018 outbreak in Mombasa, Kenya and ZIKV, MR766 strain, were used in the study. The viruses were obtained from KEMRI’s, Sample Management and Receiving Facility (SMRF).

The viruses were amplified by inoculating 200µl of each virus into individual T-25 cell culture flask (Corning Incorporated, USA) containing 70% to 80% confluent Vero cells (ATTC CCL-81). The inoculation was incubated for 1 hour at 37°C and overlayed with 5ml of Maintenance medium containing 1X Minimum essential media (MEM), (GIBCO), supplemented with earl salt, 2% of 200Mm L-glutamine (Sigma-Aldrich), and 2% penicillin-streptomycin solution containing 10,000unit/ml penicillin and 10000ug/ml streptomycin (GIBCO) and 2% Fetal Bovine Serum (GIBCO, UK). The inoculated flasks were incubated in a 5% CO_2_ incubator set at 37°C and observed daily Cytopathic effect (CPE) (22). At 80% CPE, the viruses were frozen at -80°C and harvested by thawing, then spun at 12,000 rpm for 10 minutes; the supernatant was picked, transferred to several 1.5µl cryovials, and quantified using plaque assay as described in Agha e*t al.,* 2017). The virus aliquots were then stored at -80°C until ready to use.

#### Mosquito exposure to infectious blood meals

Four-day-old female *Ae. aegypti* mosquitoes deprived of sucrose solution for 24 hours were fed with an infectious blood meal consisting of one volume of sheep blood (erythrocytes) and one volume of the respective viral suspension (either ZIKV or CHIKV) for the two sets of experiments. The mosquitoes were allowed to feed on the infectious blood meal for a duration of 45 minutes using an artificial membrane feeding system (Hemotek), adjusted to maintain the temperature of the blood-virus mixture at 37°C throughout the feeding period. An aliquot of the artificial blood meal was pipetted into a 1.5ml sterile Eppendorf tube before the blood feeding process, and promptly stored at -80°C (42) for subsequent titration using the Plague assay (33). At the end of the 45-minute feeding period, fully engorged mosquitoes were transferred to a clean BugDorm cage and maintained under controlled laboratory conditions (28 ± 1°C, 12:12 hour light-dark cycle, and 75 ± 5% relative humidity) with constant access to 10% sucrose solution (42).

#### Susceptibility and dissemination assays

A sample of the study test mosquitoes that were incubated after exposure to infectious blood meals were tested on the 7^th^, 14^th^, and 21^st^ days post-infection (dpi) for ZIKV and on the 5^th^, 10^th^ and 14^th^ dpi CHIKV. The sampled mosquitoes were then anaesthetized by placing them in a -20 freezer for 20 seconds and individual mosquitoes decapitated, by removal and transfer of their legs into a clean 1.5ml Eppendorf tube containing 500µl of homogenization media (HM). Mosquito bodies were then immobilized on a sticky surface, and a capillary tube containing HM inserted into their proboscis to stimulate salivation. The mosquitoes were left immobilized for 30 minutes, after which the capillary tubes were removed, and their content dispensed into a well-labeled 1.5ml Eppendorf tube containing 200µl of HM. The mosquito body was then picked using sterile forceps and placed in a different tube also containing 500ml of HM. The Eppendorf tube containing mosquito legs, body, and saliva were stored at -80°C awaiting cell culture assay.

##### Cell culture and inoculation

Individual mosquito bodies stored at -80°C above were thawed in ice and homogenized using 4.5mm copper beads in an Omni Bead Rapture 24 (OMNI International) set to run at the speed of 2.10mm for 20 seconds. Following the homogenization step, the homogenates were subjected to centrifugation at 12000 rpm for 10 minutes and the resulting supernatant carefully isolated and transferred into a clean cryogenic vial, ensuring the exclusion of any solid particles. Subsequently, the supernatants were used to inoculate 24-well plates containing ATTC CCL-81 Vero cells that had reached a confluence of 70-80%. For each well, 50μl of the supernatant was added and plates incubated in a 5% CO_2_ incubator for 1 hour with a 15-minute rocking interval to allow for adsorption. Next, 1ml of cell maintenance media (1X MEM [GIBCO], supplemented with earl salt, 2% of 200Mm L-glutamine (Sigma-Aldrich), and 2% penicillin-streptomycin solution containing 10000unit/ml penicillin and 10000ug/ml streptomycin (GIBCO) and 2% FBS (GIBCO, UK)) was added and the plates returned to the 5% CO2 incubator. The cells were observed daily under a microscope for the development of cytopathic effect (CPE), as an indicator of infection. Wells that showed 80% CPE were considered positive of the virus used to infect the mosquito, and the mosquitoes used to inoculate the cells deemed susceptible to the virus used to infect them. Next the legs whose bodies showed CPEs were homogenized, and the above procedure repeated. Mosquito legs that showed 80% CPE were considered to have disseminated the virus used to infect them (33,37). The same process except for the homogenization step was done on saliva samples of mosquitoes whose legs showed CPE and mosquitoes whose saliva showed CPE were considered to have transmission capabilities.

### Data management and analysis

The study data were meticulously collected following the aforementioned procedures. The genetic variability data were systematically organized and stored in a well-labeled folder to ensure easy retrieval and traceability. Similarly, the vector competence data were accurately entered into an Excel file and securely stored in a separate folder. These datasets were then subjected to rigorous analysis as outlined below, and the outcomes are comprehensively presented in this paper.

#### Genetic variability data

The raw sequences were initially edited and converted into FASTA format sequences using Chromas v2.6.6 (Technelysium). Next, Basic Local Alignment Search Tool (BLAST) was used to compare the trimmed sequences with available sequences in the gene bank. Once similar sequences were identified, the sequences were aligned using the Muscle software integrated in MEGA7. Subsequently, the sequences were sorted into the two study populations for analysis (Kisumu and Busia). The nucleotide variation of our sample population sequences were then determined using the DnaSP v6.12. This analysis helped to establish the number of haplotypes (H), haplotype diversity (Hd), and nucleotide diversity (π) in each population.

Pairwise differences and population structures were evaluated by analysis of molecular variance (AMOVA) in Arlequin 3.5.2.2, and significance was evaluated based on 10,000 permutations. The gene flow between the two localities was estimated from pairwise FST and Nm values using the DnaSP software program. For phylogenetic analysis, sequences belonging to *Ae. aegypti aegypti* (AF390098, AY432106) and *Ae. Aegypti formosus* (AY056597) was downloaded from GenBank and combined with the sequences from this study. *Ae. albopictus* (MF148303) sequence was also included as an outgroup. The combined sequences were aligned as described earlier, and the alignment was used to infer a maximum likelihood phylogeny. Phylogenetic analysis was carried out in IQTREE (46) with simultaneous evaluation of the best model and tree inference being performed based on 1000 bootstrap replicates. The phylogenetic tree was visualized in FigTree v1.4.4(47).

#### Vector competence data

The Standard software package (***Stata***, version. 15.0; ***StataCorp***) was used to determine rates of infection (The proportion of mosquito bodies that were infected with CHIKV or ZIKV), dissemination (Proportion of infected mosquitoes whose legs were positive for CHIKV or ZIKV) and transmission (The proportion of infected mosquitoes whose bodies, legs and saliva were positive for CHIKV and ZIKV) in *Ae.aegypti* at day 5, 10 and 14 post-infection (95% CI) for CHIKV and day 7, 14 and 21 for ZIKV. The same package was used to conduct a proportion test of differences to determine the statistical significance of infection, dissemination and transmission rates between the mosquito populations challenged with CHIKV and ZIKV from the two study sites.

## Acknowledgement

We gratefully acknowledge the Director General, KEMRI for providing us with the platform to conduct this study and the National Research Fund (NRF) Kenya which funded this study. We would also like to acknowledge the valuable contribution of Dr. Lihana Raphael in effectively managing project timelines and progress, which greatly contributed to the successful completion of the study within the designated timeframe.

